# Resource requirements for ecotoxicity testing: A comparison of traditional and new approach methods

**DOI:** 10.1101/2022.02.24.481630

**Authors:** Krittika Mittal, Doug Crump, Jessica A. Head, Markus Hecker, Gordon Hickey, Steve Maguire, Natacha Hogan, Jianguo Xia, Niladri Basu

## Abstract

Toxicity testing is under transformation as it aims to harness the potential of New Approach Methods (NAMs) as alternative test methods that may be less resource intensive (i.e., fewer animals, cheaper costs, quicker assays) than traditional approaches while also providing more data and information. While many stakeholders are of the opinion that this unfolding transformation holds significant promise as a more efficient and ethical way forward, few studies have compared the resources required for NAMs versus those needed for traditional animal-based toxicity tests, particularly in the field of ecotoxicology. The objective was to compare resources needed for traditional animal-based ecotoxicity tests versus alternative tests using emergent NAMs. From a bibliometric review, we estimate that traditional tests for a single chemical cost $118,000 USD, require 135 animals, and take 8 weeks. In comparison, alternative tests cost $2,600, require 20 animals (or none), and take up to 4 weeks to test 16 (to potentially hundreds of) chemicals. Based on our analysis we conclude that NAMs in ecotoxicology can be more advantageous than traditional methods in terms of resources required (i.e., monetary costs, number of animals needed, and testing times). We note, however, that the evidence underpinning these conclusions is relatively sparse. Moving forward, groups developing and applying NAMs should provide more detailed accounts of the resources required. In addition, there is also a need for carefully designed case studies that demonstrate the domain of applicability of NAMs (and make comparisons to traditional tests) to ultimately build confidence among the user community.

## INTRODUCTION

Scientific research is increasingly emphasizing the global threat posed by potential chemical contamination (Rockström et al. 2009; Landrigan et al. 2017). The traditional approach to toxicity testing of environmental chemicals and complex mixtures, which uses live animals and characterizes apical measures (e.g., survival, growth, development), has been the mainstay of toxicity testing since the 1920s. However, this approach is now shifting towards a diverse range of new approach methods (NAMs; Supplemental Figure SF1). The term NAM first originated at a European Chemicals Agency (ECHA) workshop in 2016. NAM is defined by the U.S. Environment Protection Agency (US EPA) as *any technology, methodology, approach, or combination thereof that can be used to provide information on chemical hazard and risk assessment that avoids the use of intact animals*, and thus NAMs are recognized as encompassing *any alternative test methods or strategies to reduce, refine, or replace vertebrate animals* including *in silico modeling, in vitro bioassays, early-life stage testing and toxicogenomics* (European Chemicals Agency 2016; US EPA 2019). The contemporary basis for this shift was spurred by the U.S. National Research Council (NRC) report “Toxicity Testing in the 21^st^ Century – a Vision and Strategy” (NRC 2007) which advocated modernization of toxicity testing into a more predictive, mechanistic and resource-efficient approach, and ultimately one that could better satisfy regulatory and societal needs.

Many stakeholders within academia, government, non-governmental organizations, and industry are of the opinion that the unfolding transformation in toxicity testing holds significant promise as a more efficient and ethical way forward. However, few studies have compared the resources required for NAMs versus those needed for traditional animal-based toxicity tests. Of the comparisons made, most are from the human health perspective (e.g., mammalian toxicology), and relatively little is known about vertebrate tests that underpin ecotoxicity assessments (Rovida and Hartung 2009; Settivari et al. 2015; Stanton and Kruszewski 2016; Meigs et al. 2018). The objective of this study was to compare three key resource parameters (monetary costs, number of animals needed, and time required to perform the tests) between traditional ecotoxicity testing methods that use vertebrate models against possible replacement NAMs. Analyses such as these are difficult to perform accurately due to complex testing requirements, varied national regulations, difference in type of information obtained from the two types of methods and various toxicological endpoints to be considered (Burden et al. 2016; Lillicrap et al. 2016).

Therefore, two strategies were adopted here to help overcome this difficulty. First, the objective was addressed through a mixed-methods literature review. Second, in addition to a general comparison of the three resource parameters across various ecotoxicity tests, we present a specific case, the fish acute toxicity test, for which a fish embryo test and a cell line assay have been standardized as NAMs. An evaluation of the resources required for traditional methods versus NAMs is timely and necessary to help document the extent to which emerging NAMs in ecotoxicology might indeed be more efficient, as there remain professional and organizational barriers towards the transition (Mondou et al. 2020; Pain et al. 2020).

## METHODS

We compiled data following bibliometric literature searches of specific search terms (Supplemental Table ST1). From the papers retrieved, a snowball sampling approach was taken to identify additional information sources. Only papers that provided specific numbers pertaining to the aforementioned three resource parameters were included. The bibliometric searches were conducted in October 2018 and resulted in over 1,000 publications. We focused on reports that examined standardized tests, outlined large-scale projects, and/or presented numbers for regulatory purposes. For monetary costs of most traditional tests, we relied on information from an OECD guidance document on chemical testing from 2017 and a number of publications (rows 3 to 6, Supplementary Table ST2). For the number of animals and testing times we predominantly used OECD and US EPA guidelines for various tests (rows 13 to 27, Supplementary Table ST2). As our bibliometric search yielded a relatively small database to work from, we were not able to find numbers for some tests. In these cases, we also consulted with experts in the field, obtained cost estimates via personal communication from contract testing organizations, and drew from our own experiences in conducting ecotoxicity tests. Ultimately, we gathered data from 14 OECD and 2 US EPA guidelines, 16 publications, reviews, and annual reports on animal testing, approximate costs from companies and our own experiences (Supplementary Table ST2).

Fish, birds and amphibians are the most common vertebrates used in ecotoxicology for effluent testing and testing of individual chemicals, though fish are used in the highest numbers and global estimates of fish used for effluent testing exceed 5 million per annum (Burden et al. 2016; Norberg-King et al. 2018). Further, a study examining various toxicological endpoints required for regulatory testing identified four endpoints with high fish usage where substantial savings could be realized from the incorporation of NAMs, namely, assessment of 1) acute toxicity, 2) chronic toxicity, 3) bioaccumulation, and 4) endocrine disruptors (Burden et al. 2016). Therefore, we examined resource needs for common ecotoxicity tests, across the three species, but focus on the fish acute toxicity test for the specific case study.

## RESOURCE COMPARISONS BETWEEN TRADITIONAL AND NEW APPROACH METHODS

Here, we first provide a general comparison of resources required for traditional tests and NAMs for fish, birds, and amphibians. Following this, we provide a comparison of the resources required for the specific traditional test for fish acute toxicity and the corresponding fit-for-purpose NAMs that have been proposed or accepted as alternatives.

### General comparison

We identified a total of 12 traditional tests (fish = 7; avian = 3; frogs = 2) and 7 tests using NAMs (fish = 3; avian = 3; species agnostic in vitro tests = 1). For each of these tests we examined the requirements with respect to: A) monetary costs, B) number of animals used, and C) testing times. For specific details on estimates, assumptions, and references see Supplementary Table ST3.

### Monetary Costs

The monetary costs per chemical (in USD) of common ecotoxicity tests following standard guideline tests using vertebrate models range from $15,598 for the fish acute toxicity test to $580,000 for a multi-generational test (Bottini and Hartung 2009; Willett et al. 2011; OECD 2017). Tests using alternative assays range from just under $1,000 for a primary avian hepatocyte test to $136,410 for an early-life stage fish test (Figure 1A). The median value of traditional tests ($118,000) is about 45-fold higher than the median value of alternative tests ($2,600).

**Figure 1:**
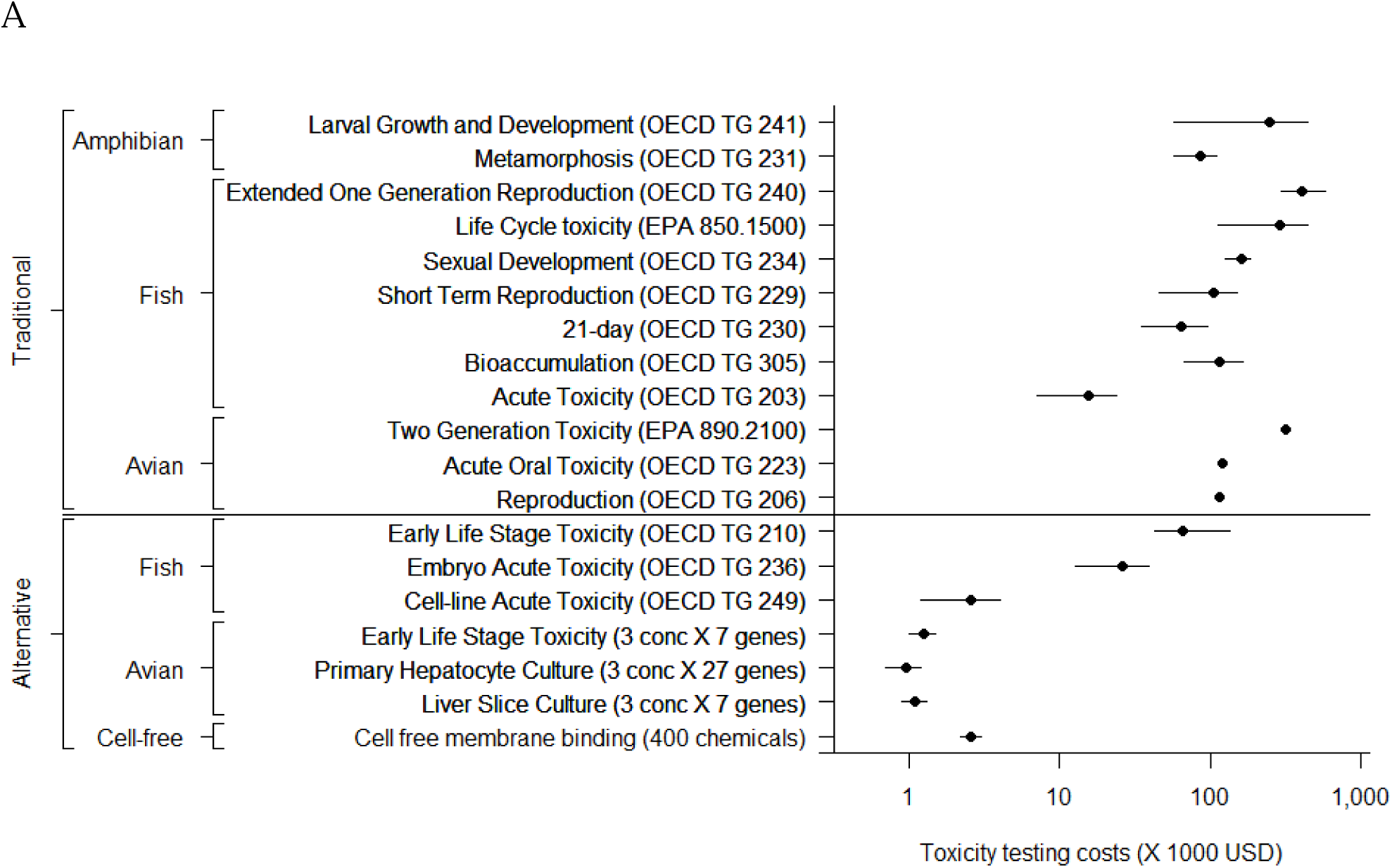
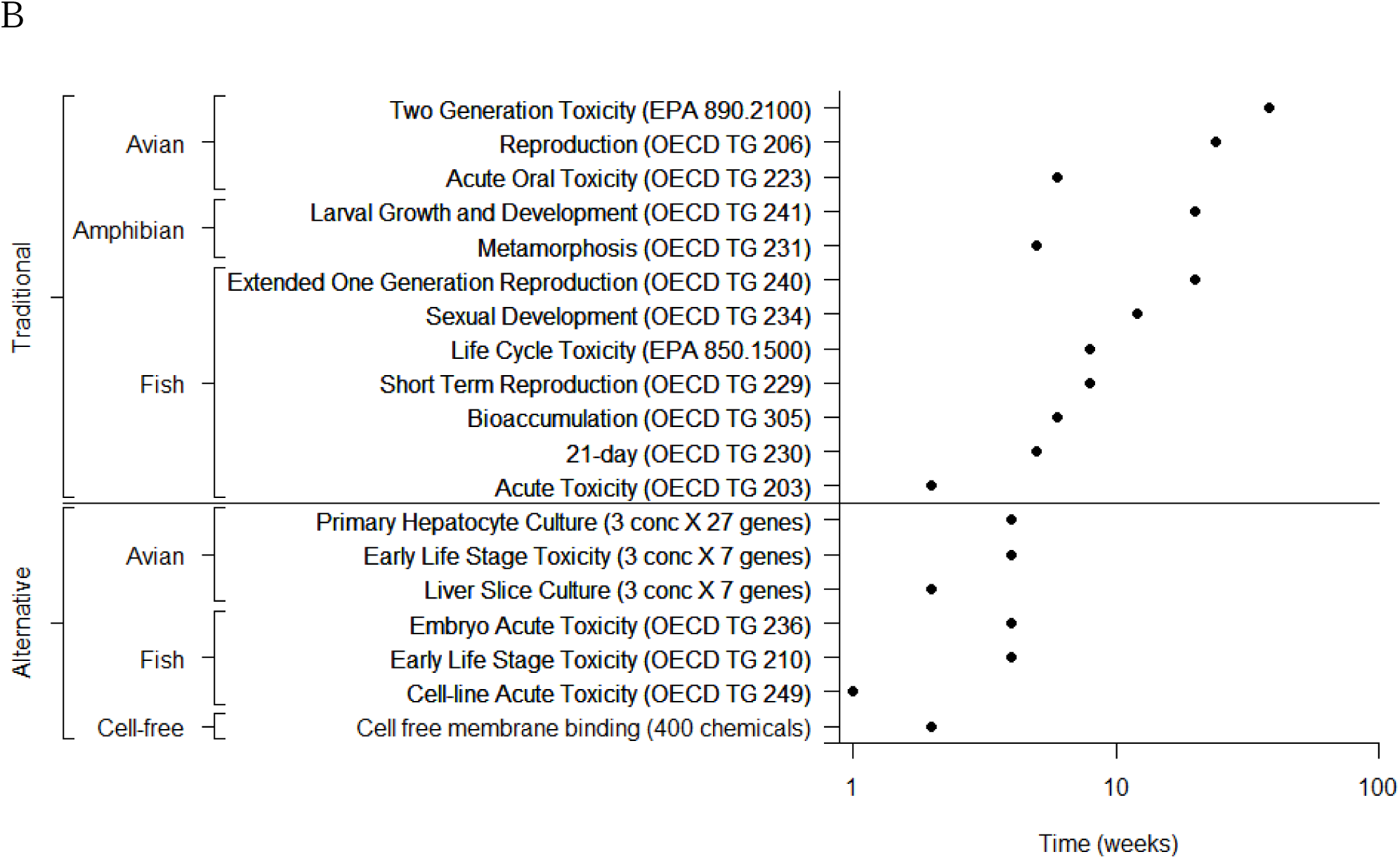
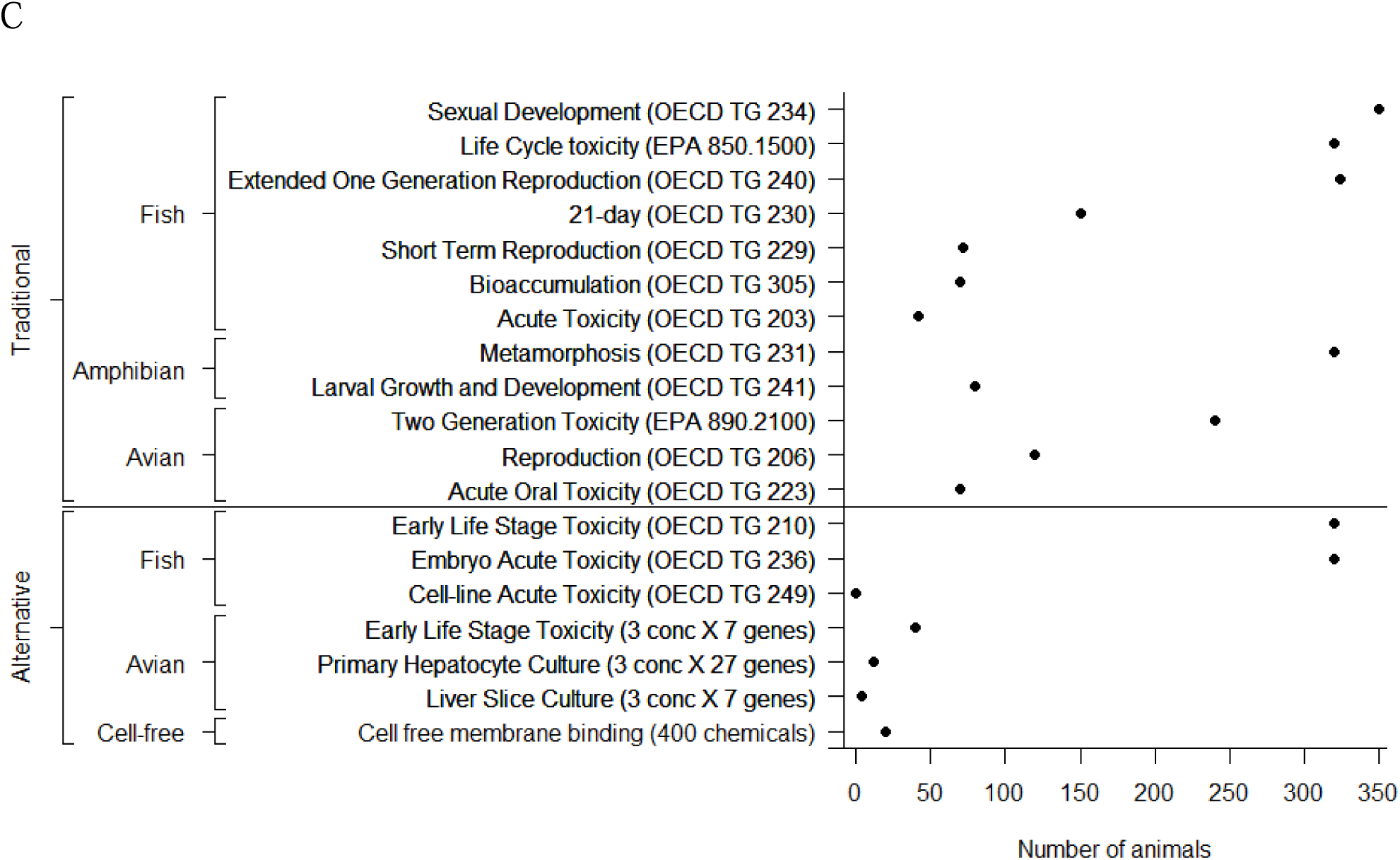
Costs associated with traditional and alternative testing strategies in terms of A) monetary costs in USD (United States Dollar; where possible, data are presented as median (black circle) and the range (bar represents minimum to maximum cost)), B) number of adult animals or embryos, and C) time in weeks. For further details on the tests and references please see Supplementary Table ST3.

The monetary costs of tests can differ across species but also within a particular species group depending on the nature of the test. Within traditional tests, we see that overall the median cost for fish varies widely from $15,598 to $411,800 per test (26-fold) whereas the cost for birds ranges from $116,000 to $319,000 per test (2.8-fold) and in amphibians ranges from $87,000 to $250,560 per test (2.9-fold). Similarly, in terms of NAMs, the cost of testing across the three fish tests ranges from $1,200 to $136,410 (113-fold), whereas the tests in birds ($700 to $1,500) and species-agnostic cell-free assays ($1,200 to $4,000) do not vary largely. The wide range in costs of the fish NAMs is largely due to the tests involving embryo exposures which require a more elaborate setup. As a point of comparison, the cost for OECD approved in vitro alternatives for eye irritation and skin sensitization tests, such as TG 430, 431, 437 and 439, range from $1,400 to $4,060 (Humane Society International; Costin 2014)

Rovida and Hartung (Rovida and Hartung 2009) estimated the monetary costs associated with chemical testing in the EU based on REACH requirements. They estimated that the number of new chemicals expected to fall under REACH would range from 68,000 to 100,000, and, using the lower estimate, they determined that monetary costs would total $13.6 billion. This scenario looked at a total of 28 tests of which the ecotoxicity tests included one avian test (OECD TG 223) and three fish tests (OECD TG 203, 210 and 305). Based on differing test requirements for specific chemicals based on their production volumes, it was estimated that 9,000 ecotoxicity tests would be needed at an estimated cost of $186 million (1.4% of the total). It is very difficult to determine how much is spent annually on ecotoxicity testing worldwide, but we propose a simple calculation, as follows. Worldwide, it has been estimated that $2.8 billion is spent annually on animal experimentation for toxicological research (Hartung 2009). Using the aforementioned 1.4% as an estimate of the share of the total expenditure realized by these four fish and avian tests, we estimate that over $39 million is spent worldwide every year for these ecotoxicity tests. This estimate, calculated based on data available from 2009, is highly simplified and likely a gross under-estimation of the true costs (e.g., it does not consider other ecotoxicity tests and study species including invertebrates, testing for environmental monitoring or compliance efforts).

### Animal Numbers

The number of animals required for standardized guideline ecotoxicity tests on vertebrate models ranges from 42 to 350 per test, while tests using NAMs call for 0 (in the case of commercial cell-lines) to 20 animals or 12 to 320 embryos per test (Figure 1B). The median number of animals needed for traditional tests (135) is over 6-fold higher than the median number of embryos or animals needed for alternative tests (20), although it is difficult to definitively make such a comparison since many alternative tests do not rely on animals (Willett et al. 2011; OECD 2017).

Within traditional tests we report that overall the median number of fish (150) and birds (120) required per test is somewhat lower than the median number of amphibians (200). In terms of NAMs, the numbers needed for fish embryos (320), avian embryos (4 to 40) and cell-based or cell-free assays (0 to 20) are higher than for rodents and other mammalian species; however, since many such tests rely on cell lines or in silico methods, the animal usage is essentially zero irrespective of the species.

Similar to our understanding of the monetary costs, there have been few estimations of the number of animals needed for traditional toxicity tests on a global scale. One paper estimated that approximately 54 million vertebrates would be needed to test about 68,000 chemicals in the EU based on REACH requirements (Rovida and Hartung 2009). Under this scenario, the number of fish and birds required for 9,000 of the four ecotoxicity tests (described in the previous section; OECD TG 223, 203, 210, and 305) was estimated at 1 million (2.2%) (Rovida and Hartung 2009). Worldwide annual estimates of the usage of animals in the laboratory range from 20 to 100 million (Taylor et al. 2008; Lush 2014). However, for countries such as the USA, these estimates do not include fish and birds and hence may be an underestimate of the true numbers of animals from these taxa. Certain countries do report the number of birds and fish used; for example, in the United Kingdom, 34,700 fish and 17,700 birds were used in 2012 for toxicology experiments; and in New Zealand 27,949 fish and over 12,000 birds were used for research, testing and teaching (Lush 2014). While it is not always possible to obtain accurate estimates for the number of birds and aquatic species used specifically for toxicity testing, using the aforementioned 2.2% as the share of birds and fish being used for the four ecotoxicity tests and depending on the estimate of total number of animals being considered (20 to 100 million), at a minimum we may estimate that the worldwide annual usage of fish and birds ranges from 440,000 to 2.2 million.

The numbers of animals being used in toxicity testing are much greater when considering environmental monitoring and regulatory compliance needs. For example, in 2018, numbers from the Canadian Council on Animal Care (CCAC) showed that in total 52,018 birds (chicken) and fish (fathead minnow, rainbow trout and zebrafish) were used for regulatory testing of products for the protection of humans, animals or the environment (CCAC 2018a). This report also stated that over 84,000 rainbow trout are used annually in relation to two key Canadian regulations governing effluent testing for metal mining and pulp and paper mill industries (CCAC 2018b). As of 2017, the compliance rate for these two regulations was greater than 97%, essentially indicating that only 3% of these effluents displayed adverse effects in the fish (i.e., only 2,520 fish exhibited symptoms) (ECCC 2017a; ECCC 2017b). In the private sector, Shell reported that in Canada, the USA, and the European Union, they used approximately 85,000 fish for regulatory testing in 2015 although this number reduced to approximately 34,000 in 2017 (Shell 2017). Looking at these numbers it appears that incorporating NAMs into monitoring and compliance testing could provide an important avenue for notable reductions in the number of animals required for monitoring and compliance purposes.

### Testing Times

The time required to conduct standard guideline tests on vertebrate models ranges from 2 to 38 weeks, while tests using NAMs typically need 2 to 4 weeks (Figure 1C). The median number of weeks needed for traditional tests (8) is about 2-fold higher than the median number of weeks needed for alternative tests (4).

Within traditional tests we see that the median number of weeks needed for avian tests (24) is 2-to 3-fold higher than the median number of weeks needed for tests involving fish (8) or amphibians (12.5). In terms of NAMs, the fish studies we report upon need 4 weeks, the avian studies need 2 to 4 weeks, and cell-free or cell-line based tests need 2 to 3 weeks. In terms of biomedical species, similar timelines (i.e., few days to a week) are expected owing to the availability of established cell lines and other alternative testing models.

There is limited information on the time needed to test panels of chemicals as would be necessary in a screening program. In the human health domain, it took an estimated 30 years to obtain toxicity data on 300 chemicals using animal tests compared to the U.S. ToxCast program which generated data across 600 mechanistic endpoints for 300 chemicals in about five years (Groff et al. 2014). Looking at the avian ToxChip as an example in ecotoxicology, transcriptomics data for 16 flame retardants using a chicken hepatocyte culture model were collected in under 4 weeks (Porter et al. 2014). In comparison, performing egg injection studies for all 16 chemicals (even if exposures were performed for two chemicals at a time) would have taken approximately 8 months to complete, and much longer (potentially years) if these were whole animal studies.

### Resource comparison of a fit for purpose test

Assessment of acute toxicity is required by REACH for registration of chemicals produced at ≥10 tons/y and is often a primary component for effluent compliance testing. The accepted traditional test for fish acute toxicity is OECD TG 203 in which small fish are exposed to test substances for a period up to 96 hrs during which lethality is monitored (OECD 2019).

Alternative tests proposed for OECD TG 203 include the fish embryotoxicity test (FET) and fish cell cytotoxicity assays. In Europe, according to the Scientific Directive on Animal Experimentation, protection is afforded to fish at the onset of exogenous feeding (Embry et al. 2010; Halder et al. 2010) and hence, many countries have adopted the FET as an alternative test. In 2013, the OECD approved the FET (i.e., Test No 236: Fish Embryo Acute Toxicity Test) as a standardized test for fish acute toxicity (OECD 2013a). In 2021, the OECD approved the rainbow trout gill cytotoxicity assay (i.e., Test No 249: Fish Cell Line Acute Toxicity) as a standardized in vitro test to predict fish acute toxicity (OECD 2021).

Resources required for the OECD TG 203 test include monetary costs that range from $7,056 to $24,140, and 42 fish per test chemical. In comparison, the FET (OECD TG 236) costs ∼$26,000 and requires 320 embryos, and the OECD TG 249 assay costs ∼$2,600, and requires no live fish since an established cell line is used. Based on the number of chemicals registered for specific productions volumes, there are an estimated 7,656 chemicals produced at over 10 tons/y (https://echa.europa.eu/reach-registrations-since-2008). Thus, conducting OECD TG 203 on all these chemicals would cost ∼$119.4 million and use ∼321,000 fish. In an alternate scenario, all 7,656 chemicals could be initially screened using OECD TG 249 (Fish Cell Line Acute Toxicity) at a cost of ∼$19.9 million and no use of animals. Assuming that 15% of these chemicals (i.e., 1,148) are flagged for subsequent toxicity testing using OECD TG 203, then the cost would be an additional ∼$17.9 million. The combined cost of this tiered approach would be ∼$37.8 million and use 48,216 fish. The overall savings would be ∼$81.6 million and ∼273,000 fish lives (Table 1).

**Table 1:** Comparison of the traditional fish acute toxicity test (OECD TG 203) and fish new approach method (OECD TG 249) to test 7,656 chemicals produced over 10 tons/y. Assume 15% of the chemicals (1,148) are prioritized for further testing. OECD = Organization for Economic Cooperation and Development; TG = Test Guideline

| Test type | Traditional | Alternative | Strategy 1 - traditional | Strategy 2 - alternative |  |  |  |
| --- | --- | --- | --- | --- | --- | --- | --- |
|  |  |  |  | OECD 249 for | OECD 203 |  |  |
|  |  |  | OECD 203 for 7,656 | 7,565 | for 1,148 |  |  |
| Test | OECD TG 203 | OECD TG 249 | chemicals | chemicals | chemicals | Total | Savings |
| Monetary cost | 15,598 | 2,600 | \$119,418,288 | \$19,905,600 | \$17,906,504 | \$37,812,104 | \$81,606,184 |
| Number of animals | 42 | 0 | 321,552 | 0 | 48,216 | 48,216 | 273,336 |
| Time (weeks) | 2 | 1 | 15,312 | 7,656 | 2,296 | 9,952 | 5,360 |

## DISCUSSION

In ecotoxicology there is a shift underway from toxicity tests that expose whole animals and measure apical outcomes to ones that use NAMs to test chemicals *in vitro* and in early-life stage organisms and yield mechanistic information. While such NAMs are considered to be cheaper, faster, and more ethical than the traditional methods, there has been a lack of empirical evidence to support such assertations. Here we aimed to synthesize information from available data-streams to provide a glimpse of the evolving field of ecotoxicity testing and the various costs associated with traditional and alternative toxicity testing methods. Such an examination is especially needed as there remain professional and organizational barriers towards this transition (e.g., concerns over error costs and pattern of familiarity (Mondou et al. 2020)).

Our analysis provides evidence that NAMs are faster, cheaper and use less animals than traditional toxicity testing methods. In terms of testing a single chemical using traditional animal tests, we estimate that the median cost of a test is $118,000 and that it requires approximately 135 animals and 8 weeks. In comparison, the median cost of an alternative test is $2,600 and would require approximately 20 animals (or 40 embryos) and up to 4 weeks to test from 16 to 400 chemicals since several chemicals may be batch-tested. Refer to Table 2 for a snapshot of the monetary cost, animals and time needed for a representative traditional and alternative test using fish (fathead minnow or zebrafish) and birds (Japanese quail).

**Table 2:**
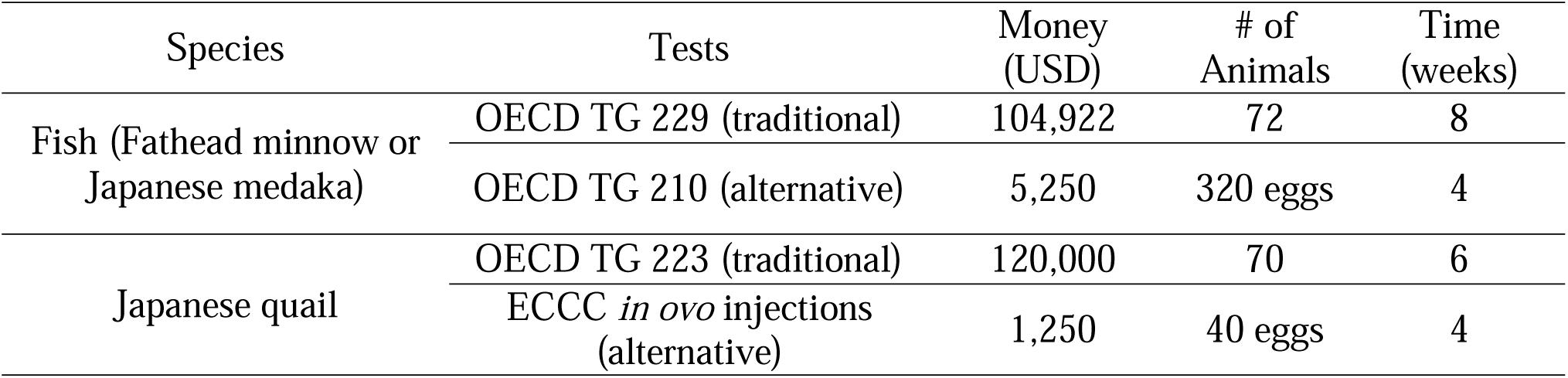
Comparison of resources needed to evaluate one chemical in a traditional (whole animal) test versus an alternative (new approach method) test for a fish and a bird in terms of monetary costs, number of animals used, and test duration. OECD = Organization for Economic Cooperation and Development; TG = Test Guideline; ECCC = Environment and Climate Change Canada

Testing is costly. For example, countries worldwide spend 7 to 24 billion dollars annually on pollution abatement and control activities (Statistics Canada 2012; MAPI 2015; Eurostat 2017). In terms of testing chemicals, scenarios out of the European Union present numbers that extend into billions of dollars (Rovida and Hartung 2009). Even if the backlog of chemicals were adequately tested, there will always be a need to perform toxicity tests given the introduction of 500 to 1000 new chemicals annually into commerce (Arnold 2015) and the growing number of environmental sites that require monitoring. Some reports have started to investigate whether the incorporation of NAMs into testing programs may realize monetary and non-monetary cost savings. An earlier study concluded that animal testing needs could be reduced by up to 70% by the adoption of intelligent testing strategies such as quantitative structure activity relationships (QSARs), grouping and read-across methods (Van der Jagt et al. 2004). A more recent study estimated that 3 – 15% of chemicals initially screened using NAMs would be prioritized for *in vivo* testing as part of a tiered testing program (Thomas et al. 2017). These initial studies demonstrate that NAMs may help increase efficiencies though more rigorous evaluations are needed.

Incorporation of NAMs in toxicity testing is starting to be realized by key stakeholders in the human health domain. A key example is the U.S. EPA’s ToxCast program that has screened thousands of chemicals via hundreds of *in vitro* assays at a fraction of the cost to test these chemicals using animal bioassays (Dix et al. 2007; Thomas et al. 2019). In ecotoxicology, regulatory decisions still rely on the outcomes of whole animal studies though progress is being made in terms of adopting NAMs. First, we are seeing the emergence and acceptance of new testing systems that may serve as alternatives to animal tests. In 2018, the U.S. EPA listed OECD Test Guideline #236 (Fish Embryo Acute Toxicity, FET, Test) as an “*alternative test methods or strategies the Administrator has identified that do not require new vertebrate animal testing and are scientifically reliable, relevant, and capable of providing information of equivalent or better scientific reliability and quality to that which would be obtained from vertebrate animal testing”* (US EPA 2018). However, European researchers examining the ability of this OECD Test Guideline #236 to predict outcomes in standard acute fish toxicity tests for regulatory purposes highlighted several limitations (e.g., the fish embryo test does not capture key modes of action, or is challenging to use for certain classes of chemicals) (Sobanska et al. 2018), thus illustrating the need for more research activities on the fish embryo test system. For birds, researchers at Environment and Climate Change Canada (ECCC) have proposed the standardization of early-life stage toxicity tests using avian eggs though this particular study only evaluated eight chemicals in one avian species and thus there is a need for additional studies (Farhat et al. 2019).

Second, there is a need for NAMs to be made available in a more consistent and commercial manner while also being affordable and reliable. Within the ‘omics fields we are starting to see the arrival of products in the marketplace that can aid in the investigation of the transcriptome, proteome, and metabolome of species of ecotoxicological interest. For example, a Canadian team of academic, government and industry partners is co-designing 384-well qPCR arrays (EcoToxChips) and a corresponding data evaluation tool (www.ecotoxxplorer.ca) to help characterize, prioritize and manage environmental contaminants and complex mixtures of regulatory concern (Basu et al. 2019). We estimate that coupling such a toxicogenomic tool with alternative testing systems (e.g., the aforementioned fish embryo test or avian egg injection method) may enable rapid and deeper hazard characterization for ∼$1,000 −5,000 per tested chemical.

Finally, as we enter a big data era, the information resulting from NAMs must be rapidly processed and be amenable for decision-making under a range of contexts. Frameworks such as adverse outcome pathways (AOPs) and the OECD’s AOP Knowledgebase (https://aopkb.oecd.org/) along with standardized reporting templates and Findable, Accessible, Interoperable and Reusable (FAIR) principles could better help enable the user community to maximize the use of data. The ecotoxicological community is also starting to benefit from a diverse set of relevant and publicly accessible tools that allow users to efficiently query large databases of c186.926
hemicals and toxicological information (e.g., U.S. EPA’s CompTox Dashboard and the ECOTOX Knowledgebase), perform species read-across assessments (U.S. EPA’s SeqAPASS, EnviroToxDatabase.org), conduct risk assessments (HESI Risk 21, CAFÉ), derive transcriptomic points of departure (BMDExpress2, FastBMD-www.fastbmd.ca), and analyze various ‘omics data: EcoToxXplorer (www.ecotoxxplorer.ca), NetworkAnalyst (www.networkanalyst.ca), MetaboAnalyst (www.metaboanalyst.ca), and MicrobiomeAnalyst (www.microbiomeanalyst.ca).

Nevertheless, there are a number of challenges associated with the adoption of NAMs. For instance, the two approaches examined here – traditional methods and NAMs – provide different types of data; risk assessment and regulatory decisions are typically made based on apical results including mortality, reproductive or developmental effects, which are obtained from traditional methods. However, NAMs typically provide mechanistic information including cytotoxicity, receptor binding, enzyme activity, and large sets of omics data (Villeneuve and Garcia-Reyero 2011). Extrapolating such results from NAMs across various levels of biological organization, i.e., sub-cellular and cellular level to predict effects in the whole organism in an accurate manner presents a major challenge in obtaining biologically relevant information (van Vliet 2011). Thus, the predictive capacity of NAMs to whole animal methods is one of the main obstacles to implementing and integrating them into the decision making process.

The question regarding the biological relevance of data obtained from these methods leads to the issue of acceptance of the methods within regulatory bodies. While there has been increased interest in the development of these types of methods, acceptance of these methods within the ecotoxicological community is lacking. Thus, if the NAMs being developed and validated do not gain acceptance, they are perhaps of not much practical use regardless of how resource efficient they may be. Other challenges associated with NAMs are related to the difference in national regulations. Data from studies for the same endpoints in one country may not be acceptable in other countries thus resulting in the need to repeat the study and thereby increase costs. Further, the analyses of complex data generated by NAMs often require specialized skills and knowledge and thus calling upon additional assistance for data analysis can add to the total costs.

Based on our analysis, here we conclude that NAMs in ecotoxicology can be more advantageous than traditional methods in terms of resources required (i.e., monetary costs, number of animals needed, and testing times). However, there is a need for carefully designed case studies that demonstrate the domain of applicability of NAMs to ultimately build confidence among the user community (Kavlock et al. 2018). Thus, we note that the evidence underpinning these conclusions is relatively sparse and that moving ahead, groups developing and applying NAMs should provide more detailed accounts of the resources required.

