## Supplemental information for "Resource requirements for ecotoxicity testing: A comparison of traditional and new approach methods"

**SUPPORTING INFORMATION**

**Supplementary table ST1**: List of bibliometric terms searched in Web of Science (searches performed October 2018).

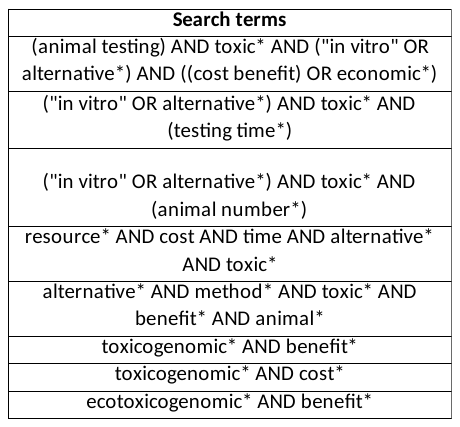

**Supplementary Table ST2:** List of publications and reports from which information pertaining to monetary costs, number of animals used, testing times or comparisons between methods was extracted.

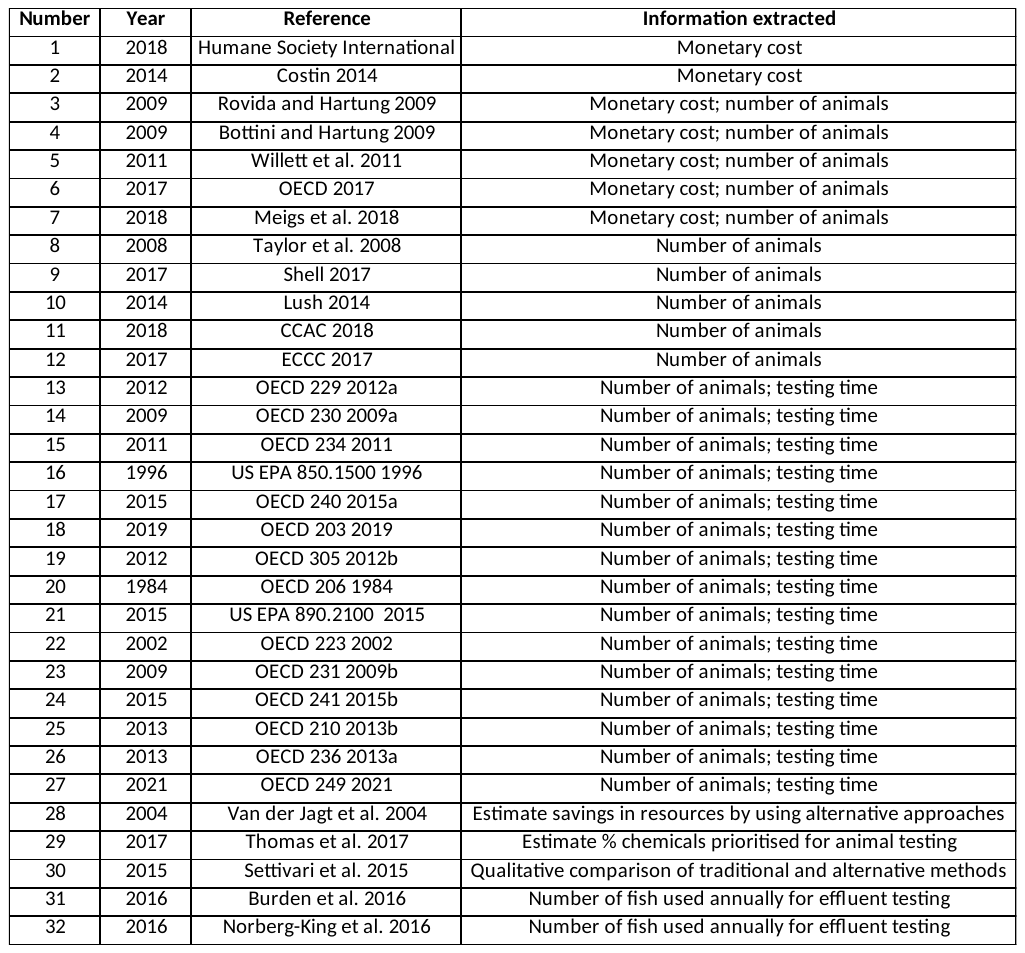

**Supplementary Table ST3**: Monetary cost (in USD), number of animals needed, and test duration (in weeks) of traditional (whole animal) tests versus alternative (new approach method) tests. OECD = Organization for Economic Cooperation and Development

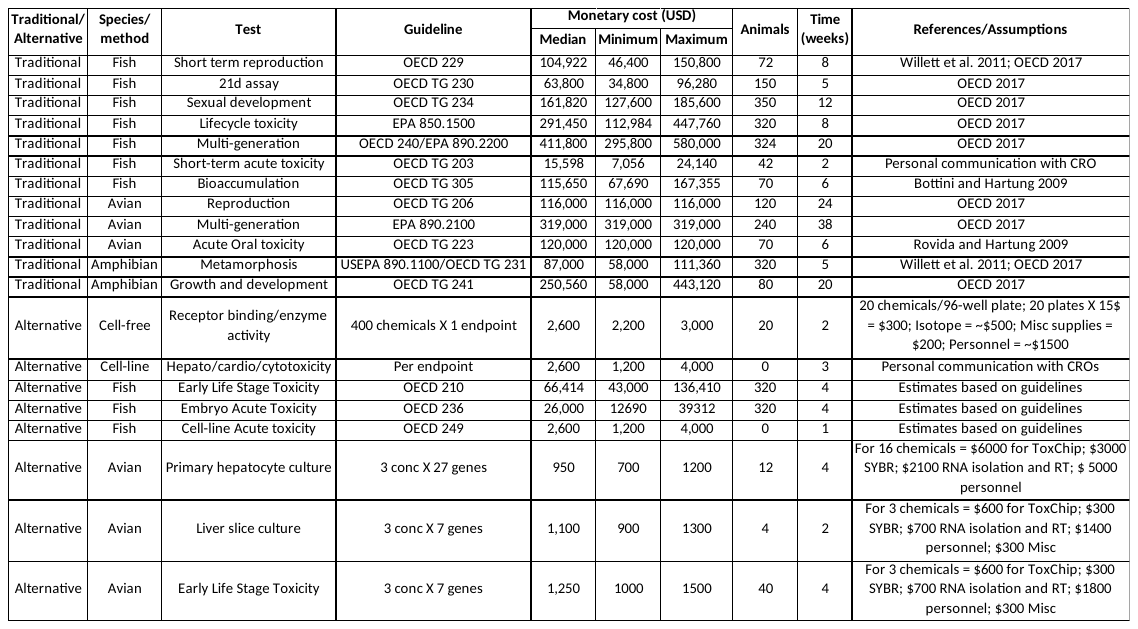

**Supplementary Figure SF1**: Graphical representation of bibliometric searches in Web of Science for the overall field of environmental toxicology and further refined with key words related to human/mammalian, fish, or avian species (searches performed in January 2020).

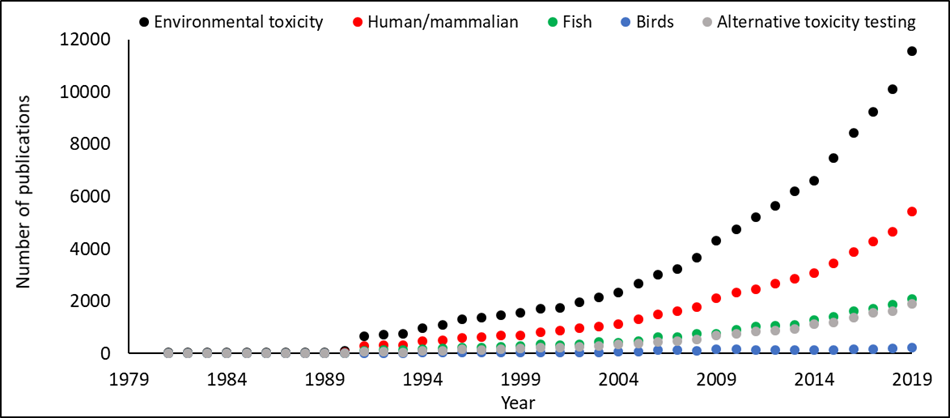
